# Recognizing the opponent: the consolidation of long-term social memory in zebrafish males

**DOI:** 10.1101/2023.02.03.527013

**Authors:** Luciano Cavallino, María Florencia Scaia, Andrea Gabriela Pozzi, María Eugenia Pedreira

**Affiliations:** Laboratorio de Neuroendocrinología y comportamiento en peces y anfibios, DBBE, IBBEA-CONICET, Facultad de Ciencias Exactas y Naturales, Universidad de Buenos Aires, CABA, Argentina. Intendente Güiraldes 2160, Pabellón 2, Piso 4°, Laboratorio26 (C1428EHA).; Instituto de Fisiología, Biología Molecular y Neurociencias (IFIBYNE), CONICET, Facultad de Ciencias Exactas y Naturales, Universidad de Buenos Aires, Buenos Aires, Argentina

**Keywords:** Aggressive behaviors, long-term social memory, NMDA receptor antagonist, zebrafish

## Abstract

Recognizing and remembering another individual in a social context could be beneficial for individual fitness. Especially in agonistic encounters, remembering an opponent and the previous fight could allow avoiding new conflicts. Considering this, we hypothesized that this type of social interaction forms a long-term recognition memory lasting several days. It has been shown that a second encounter 24 hours later between the same pair of zebrafish males is resolved with lower levels of aggression. Here, we evaluated if this behavioral change could last for longer intervals and a putative mechanism associated with memory storage: the recruitment of NMDA receptors. We found that if a pair of zebrafish males fight and 48 or 72 hours later fight again, they resolved the second encounter with lower levels of aggression. However, if immediately after the first encounter opponents were exposed to MK-801 (NMDA receptor antagonist), they resolve the second one with the same levels of aggression, that is, no reduction in aggressive behaviors was observed. These results suggest the formation of a long-term social memory related to recognizing a particular opponent and/or the outcome and features of a previous fight.

## 1. Introduction

In nature, animals live in a complex environment and their capacity to learn and remember is beneficial for their fitness. In particular, the interactions between members of their own species may result in competition for resources such as territories, food, or mates. These aggressive interactions represent valuable experiences that can modulate future behaviors. For example, if an animal engages in an aggressive agonistic encounter and loses, the next time it faces the same opponent the loser could prevent possible injuries by avoiding a new fight. This is possible in the context of the formation of a social memory in which the individual remembers their opponent and the contest outcome (Johnsson, 1997).

Memory formation and duration have been widely studied in humans and non-human animals. It is possible to distinguish between short-term memory and long-term memory (LTM), each presenting different characteristics and peculiarities (Cowan, 2008). In particular, LTM refers to those cases in which individuals can retrieve the acquired information for at least 24 hours after the learning event. It is well known that the process of LTM formation involves de-novo RNA and protein synthesis (McGaugh, 2000). Moreover, protein degradation is also involved, so both processes interact in a coordinated manner to properly form and store memories (Jarome & Helmstetter, 2014).

In some agonistic interactions, a social hierarchy or a dominance relationship is established, and this could be maintained over time (Paull *et al*., 2010). A LTM formation related to this type of interaction could allow individuals to remember and recognize their opponents, maintaining the social dynamic. For example, in fights between paradise fish, males can remember their opponent and change their behavior in a subsequent encounter (according to the previous outcome); this memory lasts for 24 hours but is undetectable after one week (Miklósi *et al*., 1992).

Learning and recalling new information and experiences represents a permanent readjustment to a changing environment, allowing individuals to change their behavior accordingly. In this context, it may be critical to maintain memories for long periods of time. The evolutionary advantage of LTM in nature is evident throughout different vertebrate groups such as fish: sharks exposed to a capture situation decrease their probability to be re-caught (Mourier *et al*., 2017), while cleaner fish show hiding behaviors eleven months after experiencing captivity (Triki & Bshary, 2020). These pieces of evidence show the importance of retaining memories for long periods of time and the direct effect it has on the survival of individuals. Furthermore, considering individuals living in a complex social environment, memories related to recognizing and remembering other individuals could lead to better resolving the agonistic interactions (Huntingford, 2013), thus, establishing a social memory of familiar conspecific for several days could be important for the stability of social groups (Kogan *et al*., 2000). Consequently, it is interesting to evaluate the duration of memories related to individual recognition and social interaction.

In general, mechanisms related to storing LTM are conserved throughout evolution. In particular, the participation of the glutamatergic system is well described (Aigner, 1995). Given the relation between long-term potentiation and the formation of LTM, different reports have demonstrated the role of N-methyl-d-aspartate (NMDA) receptors in memory consolidation. These ionotropic receptors are sensitive to glutamate and permeable to Na^+^, K^+^ and Ca^2+^. The pore of the channel is blocked by Mg^2+^, which after a depolarization, is expelled allowing a Na^+^, K^+^ and Ca^2+^ current. The depolarization and glutamate binding maximize the Ca^2+^ influx, altering the synaptic efficiency through an intracellular cascade of signaling and thus promoting the LTM formation (Luscher & malenka, 2012). As a consequence, an antagonist of this type of receptors has been used to impair LTM formation (McGaugh & Izquierdo, 2000). Dizolcipine, MK-801, is a noncompetitive NMDA receptor antagonist, which binds inside the channel preventing the Ca^2+^ influx and inhibiting the long-term potentiation (Nicoll & Malenka, 1999). Evidence in zebrafish suggests that MK-801 prevents memory formation in single-trial inhibitory avoidance (Blank *et al*., 2009), contextual fear conditioning test (Kenney *et al*., 2017), and associative learning performance (Sison & Gerlai, 2011). MK-801 can have behavioral and social effects since exposure for 15 minutes decreases time spent closer to the conspecifics zone and to the segment nearest a mirror image (Zimmermann *et al*., 2016) and, even if it does not affect the overall swimming activity, it increases specific swimming in circles (Swain *et al*., 2004). Interestingly, these effects on behavior have been observed when individuals are tested immediately after drug treatment, but no effect on locomotion is found 24 hours after individuals have been intraperitoneally injected, 2 mg/kg. (Franscescon *et al*., 2020).

Zebrafish, *Danio rerio*, is a promising species to study memory in a non-mammalian vertebrate model, due to their ability to learn and recall different tasks like inhibitory avoidance (Blank *et al*., 2009), Y-maze memory task (Cognato *et al*., 2012), and social recognition memory (Madeira & Oliveira, 2017), among others. The numerous genetic markers developed in this species, and the functional similarities between mammalian and zebrafish genes make it a suitable model to investigate memory and its mechanisms as a first step to understanding this process in other vertebrates (Gerlai, 2016). Zebrafish is a shoaling species showing conserved social behaviors, such as mating and fighting for different resources, like territory or food (Kalueff *et al*., 2013). They exhibit aggressive behaviors and hierarchical dominance (Paull *et al*., 2010), and, in this context, it is very useful to recognize and remember other individuals in order to avoid unnecessary fighting, physical damage or energy waste. In both sexes, intrasexual dyadic encounters usually start with a display (fish erects its dorsal and anal fins and flares its body flank toward the opponent) and it is followed by bites, circles (defined by two fish approaching each other in opposite directions and antiparallel position displaying circumference), and chases (when individuals actively pursuit each other alternately), until one individual wins the fight and chases the loser, which performs submissive behaviors as fleeing or freezing (Oliveira *et al*., 2011, Scaia *et al*., 2022). Previous evidence suggests that zebrafish males show the ability to recognize a particular opponent, remember fight outcomes, and change their fighting strategies in a subsequent encounter against the same opponent, even after a 24-hour interval between fights (Cavallino *et al*., 2020). After previous experience against the same opponent, fish resolves subsequent encounters with reduced levels of aggression (number of bites and total time of aggressive behaviors), but this reduction is not observed if the subsequent opponent is different, suggesting a social memory formation (Cavallino *et al*., 2020).

Our working hypothesis was that zebrafish males form a long-term memory associated with the first encounter. Therefore, the present work pursued two aims: The first one was to evaluate if the recognition and retrieval of a particular opponent, and the fight between them, last for more than 24 hours, and the second to evaluate the role of NMDA receptors on the acquisition of long-term social memory. To reach such a goal, we assessed aggressive behaviors in male-male zebrafish dyads, during successive encounters, with 48 or 72-hour intervals between fights, to evaluate longer retention than 24 hours. In another set of experiments, to evaluate one mechanism related to LTM formation, individuals were exposed to water or NMDA antagonist, MK-801, immediately after the first encounter to impair LTM consolidation.

## 2. Materials and methods

### 2.1 Ethics statement

The procedures used in this study followed the institutional guidelines for the use of animals in experimentation (Comisión Institucional para el Cuidado y Uso de Animales de Laboratorio, Facultad de Ciencias Exactas y Naturales, Universidad de Buenos Aires, Protocol 75b/2021) and were in accordance with National regulations (Comité Nacional de Ética en la Ciencia y la Tecnología). All procedures were in compliance with the Guide for Care and Use of Laboratory Animals (eighth ed. 2011, National Academy Press, Washington, p. 220.) and the ARRIVE guidelines.

### 2.2 Animals and maintenance

Zebrafish adult males (160 individuals: standard length, SL= 2.47±0.02 cm, Weight, W=0.199±0.004g) were obtained in a commercial aquarium. They were acclimated one month before the experiment in different 20-L aquaria of dechlorinated filtered water (Density: 1 fish per liter), in appropriate conditions and controlled water quality: 25°C-26°C, pH= 7.5/7.8, a photoperiod of 14h light (8am-22pm): 10h dark cycle (22pm-8am), constant aeration (Avdesh *et al*., 2012), and were fed twice daily with commercial food (Tetra®). During the test the same temperature, water and photoperiod conditions were maintained and every pair of opponents were fed once, two hours after the test, with the same amount of food.

### 2.3 Experimental protocol

Following the same protocol of Cavallino and coworkers (Cavallino *et al*., 2020), two size-matched adult zebrafish males from different community tanks were isolated in a small aquaria (2 L, 15 cm (high) x24cm (length) x9cm (width)). Individuals were separated by an opaque perforated barrier, which allows chemical but not visual communication, for 24 hours before the experiment. After this period of isolation, the barrier was removed and the encounter between males was registered with a JVC Everio video camera, for 30 minutes. Then, individuals were separated again as previously explained, and after 48 or 72 hours the barrier was removed and the individuals were exposed to the same opponent again (Figure 1a). Sixty individuals were used in these experiments (15 pairs for the 48 hours interval and 15 pair for the 72 hours interval).

**Figure 1.**
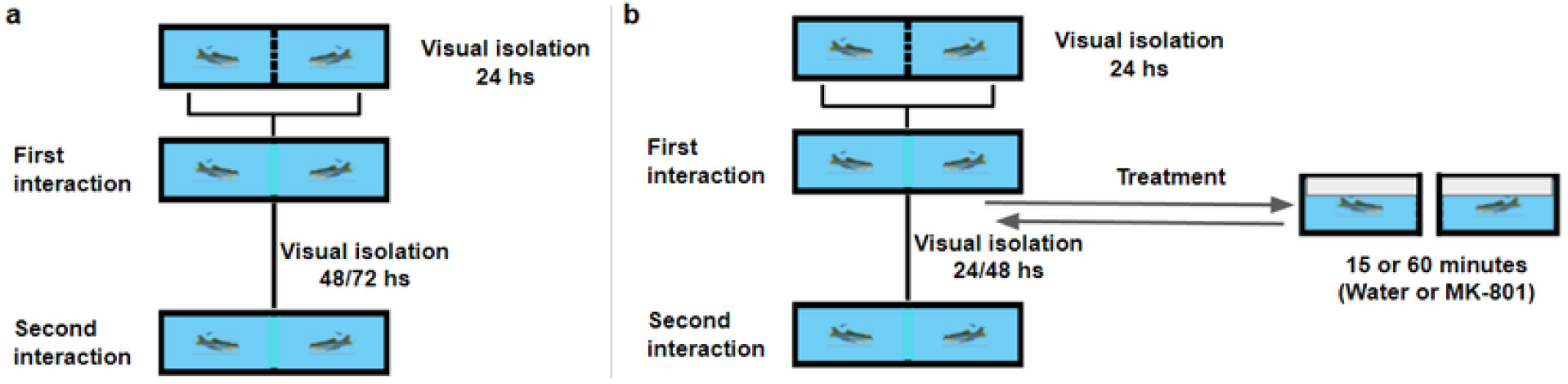
Experimental devices in which pairs of zebrafish males were isolated in 2 L aquarium, separated by a barrier. Twenty four hours later, individuals were allowed to interact for 30 minutes until separated again. a) 48 or 72 hours later the same pairs interact again for 30 minutes. b) Immediately after the first encounter, pairs were moved to separate aquaria for 15 or 60 minutes, containing water or MK-801 dissolved in water. After treatment, pairs were returned to their tank and isolated for 24 or 48 hours until they interacted again for 30 minutes.

In another set of experiments, immediately after the first encounter in a test tank, individuals were moved to a treatment aquaria (100 ml, 6cm (high) x10cm (length) x10cm (width)) containing pure water or 20μM of MK-801 dissolved in water. After a 15- or 60-minute treatment, individuals were placed again in the respective compartment of their test tank. Then, individuals were isolated as previously explained, and faced the same opponent 24 or 48 hours later (Figure 1b).

Aggressive behaviors were distinguished according to the ethogram described for male dyadic encounters of this species and included displays, circles, bites and chases (Oliveira et al., 2011). The number of bites and the total time of aggressive behaviors were measured during the fight phase, which started with the first aggressive behavior and finished with the resolution of the conflict. We measured only the time that fish exhibit aggressive behaviors as displays, chase, circle, or/and bites to determine the level of aggression of each encounter. We also quantified the number of bites, since they are the most aggressive behavior involving body contact between opponents. One hundred individuals were used in these experiment (10 pairs for each treatment)

All the experiments were performed at the same time range (10:30-13 hours) to avoid possible behavioral fluctuations due to circadian rhythm. It is worth mentioning that in pilot experiments encounters performed later (14-16 hours) showed lower (or none) levels of aggressive behaviors. No water changes were made during the experiments.

After the final encounter, individuals were measured, weighted and sacrificed to confirm the sex. Individuals were euthanized by a rapid chilling followed by decapitation (American Veterinary Medical Association, AVMA, Guidelines for the Euthanasia of Animals, 2020). All fish used in this work had been correctly identified as males.

All the experiments were conducted in a room with constant temperature and photoperiod as previously described in section 2.2, additionally, the same light intensity was maintained during the encounters. Test aquaria were surrounded by an opaque blue background, leaving the top and front side uncovered. We recorded with a JVC Everio video camera placed in front of the aquaria.

### 2.4 Drug treatment

MK-801 solution was prepared following the protocol of Blank and coworkers (Blank *et al*., 2009). Briefly, the 20 μM dilution was prepared by dissolving 0.675 mg of (+) MK-801 hydrogen maleate (Sigma-Aldrich) in 100 ml of filtered water. Fifteen minutes of 20 μM MK-801 was chosen because in previous experiments in zebrafish this treatment had an amnesic effect (Blank *et al*., 2009, Kenney *et al*., 2017). Since no such effect was observed in our experiments, we performed a second experiment using a 60 minutes treatment.

### 2.5 Statistical analysis

All analyses were done in R studio (R i386 3.6.2). Considering that data violate the assumption of parametric analysis, non-parametric or Bayesian tests were performed. Mann Whitney tests were performed to compare unpaired data and morphological traits (SL and weight) (using wilcox.test(paired=FALSE)). A Bayesian analysis was performed to compare the total time of aggressive behaviors and the number of bites between subsequent encounters (Experiment of 48 and 72 hours interval, and water (15 and 60 minutes) Mk-801 (15 and 60 minutes), and MK-801 48 hours interval (60 minutes) treatments). As every pair of individuals faced each other twice, an index for each variable was calculated as X1-X2 (e.g., number of bites in the first encounter-number of bites in the second encounter). This index was compared with a theoretical mean of 0 (No differences between encounters), and a non-informative prior. Lower Credible Interval (LCL) and Upper Credible Interval (UCL) were calculated as well the p-value with α = 0.05, one-tailed test. Data are presented as mean±SEM.

As a complement to the Bayesian analysis, a non-parametric test (Paired Wilcoxon signed rank test) was performed to compare between two paired data samples (using wilcox.test(paired=TRUE) and wilcox_test(paired=TRUE) from package “coin”). R scripts are available from the corresponding authors upon reasonable request.

## 3. Results

### 3.1 Morphological traits

Since size-matched adult males were used for dyadic agonistic encounters, morphological traits were compared and no differences were found between winners and losers in their weight (Loser W=0.188 ±0.007 g; Winner W=0.203 ±0.007 g; N=80, p=0.077) or standard length (Loser SL=2.41 ±0.03 cm; Winner SL=2.48± 0.03 cm; N=80, p=0.095).

### 3.2 Effect of 48- and 72-hours interval between encounters on aggressive behaviors

In order to evaluate if the reduction in the total time of aggressive behaviors observed in subsequent encounters (Cavallino *et al*., 2020) remains for longer than 24 hours, 48- and 72-hour intervals were tested.

In reference to the 48-hour interval, the total time of aggressive behaviors was reduced in the second encounter when compared to the first one, being the difference between days different from zero (LCL=94.47 sec, UCL=256 sec, Z=-3.56, n=15, p-value=0.00017, Fig 2a). A reduction in the number of bites was also observed in the second encounter (LCL=6.85 bites, UCL=76.41 bites, Z=-1.96, n=15 p-value=0.024, Figure 2b). In all experiments, the contest outcome was maintained in the second encounters.

**Figure 2.**
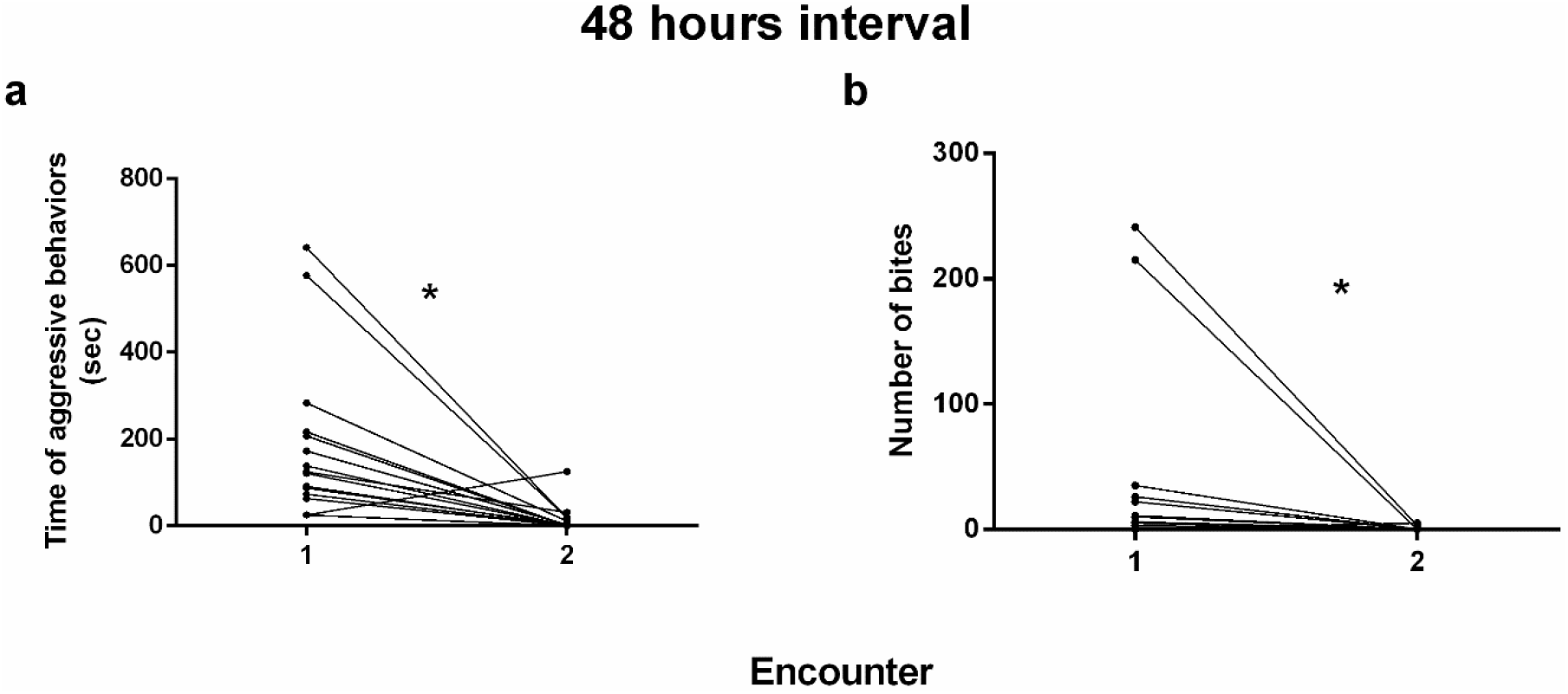
a) Total time that opponents performed aggressive behaviors (in seconds). Each line represents a pair of opponents with a 48-hour interval between encounters. (Encounter 1: 189.53±47.78 sec; Encounter 2: 13.86±8.36 sec, data is presented as mean±SEM). b) Number of bites performed during encounters. Each line represents a pair of opponents with a 48-hour interval between encounters. (Encounter 1: 42.21±21.25 bites; Encounter 2: 0.57±0.40 bites, Data is presented as mean±SEM). Asterisks (*) represent significant differences between encounters, with α=0.05.

Similarly to 48 hours interval, the total time of aggressive behaviors and number of bites were also determined for the first and second encounters, in this case with a 72-hour interval between them.

The total time of aggressive behaviors was also higher in the first encounter when compared to the second one (LCL=72.90 sec, UCL=437.03 sec, Z=-2.30, n=15, p-value=0.010, Figure 3a). Moreover, the total number of bites was also higher in the first encounter compared to the second one (LCL=48.47 bites, UCL=243.77 bites, Z=-2.46, n=15, p-value=0.006, Figure 3b). In all experiments, the contest outcome was maintained in the second encounter.

**Figure 3.**
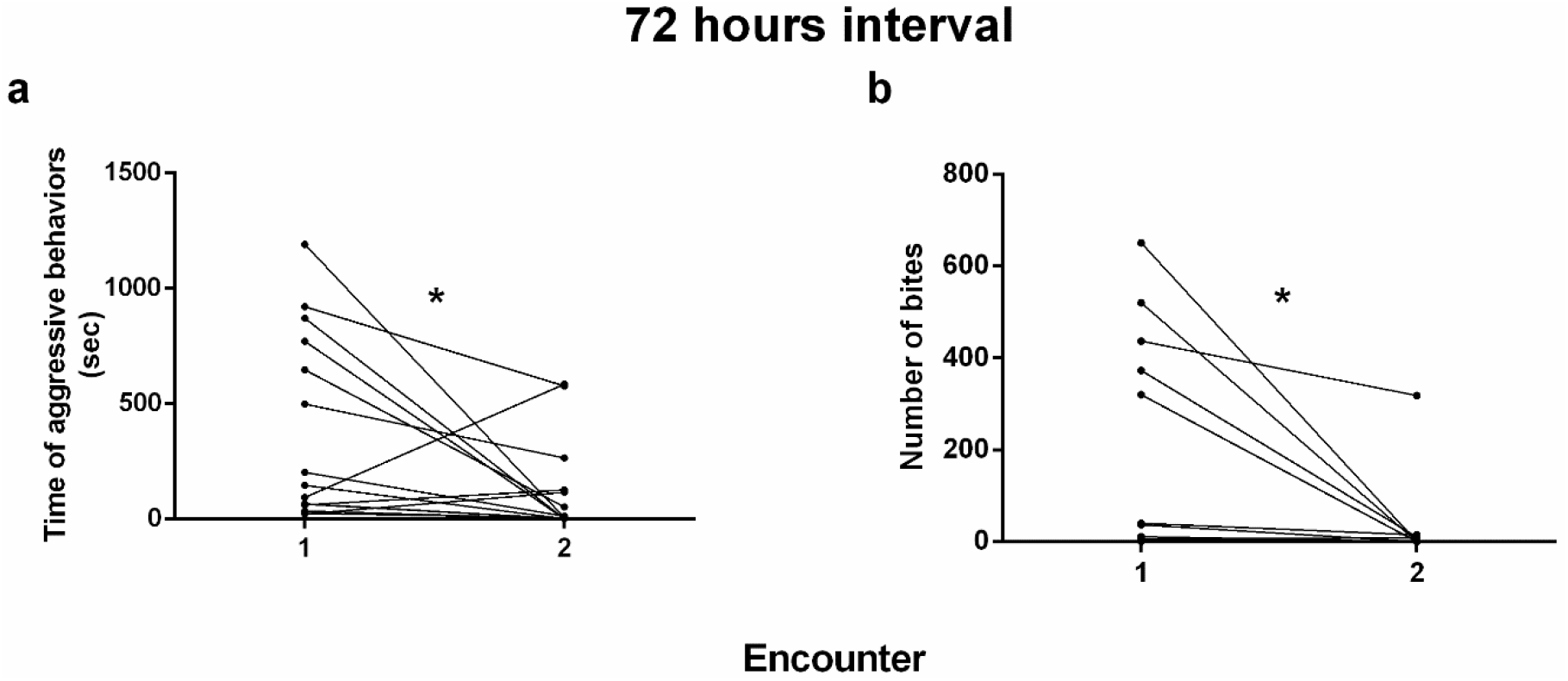
a) Total time that opponents performed aggressive behaviors (in seconds). Each line represents a pair of opponents with a 72-hour interval between encounters (Encounter 1: 373.86±104 sec; Encounter 2: 115.73±52.27 sec, data is presented as mean±SEM). b) Number of bites performed during the encounter. Each line represent a pair of opponents with a 72-hour interval between encounters. (Encounter 1: 171.78±62.75 bites; Encounter 2: 25.14±22.56 bites, Data is presented as mean±SEM). Asterisks (*) represent significant differences between encounters, with α=0.05

The effect observed in 24-hour intervals (Cavallino et al., 2020) is maintained for at least 72 hours. After reaching the first goal, we then pursued the next series of experiments. A treatment with MK-801 has shown amnesic effects in different memory tasks (Blank *et al*., 2009; Kenney *et al*., 2017). Here this treatment with such drug was performed in order to evaluate the effect of impaired long-term memory consolidation reflected by changes in the outcome and/or features of the second encounter.

### 3.3 Effect of 15 minutes MK-801 exposure in a 24-hours interval

Initially, we chose the lower time of treatment described in the literature (Blank *et al*., 2009). Thus, when individuals were exposed to water for 15 minutes immediately after the first encounter, total time of aggressive behaviors was significantly lower in the second encounter when compared to the first one, being the difference between days different from zero (LCL= 62.12 sec, UCL=249.11 sec, Z=-2.73, n=10, p-value=0.0030, Figure 4a). The same effect was observed in individuals exposed to MK-801 for 15 minutes (LCL=97.67 sec, UCL=370.48 sec, Z=-2.82, n=10, p-value=0.0023, Figure 4b). It is worth mentioning that, in parallel to the 48- and 72-hour intervals, in all the experiments the winner of the first encounter also resulted in a winner during the second one.

**Figure 4.**
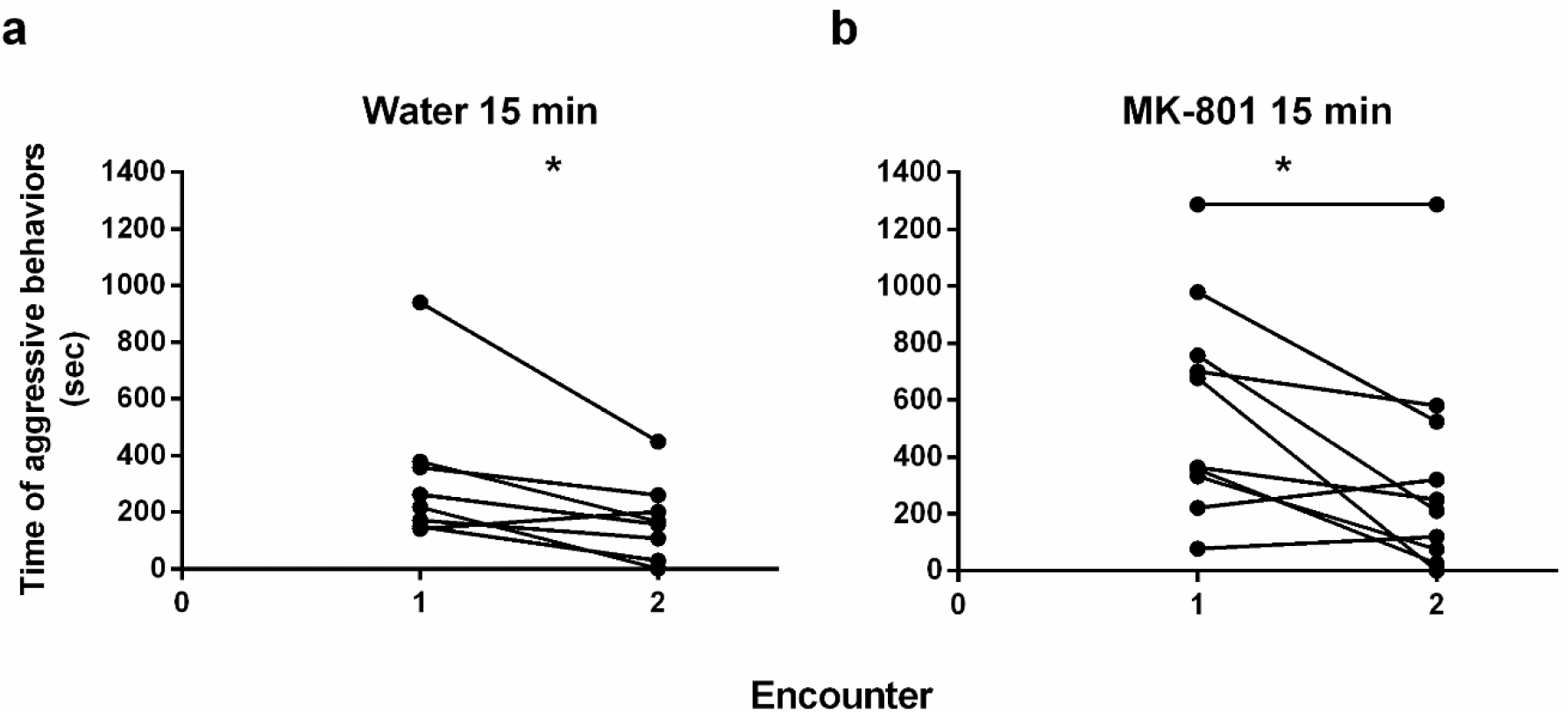
Total time that opponents performed aggressive behaviors (in seconds). Each line represents a pair of opponents with a 24-hour interval between encounters. Immediately after the first encounter individuals were exposed to: a) 15 minutes of water (Encounter 1:306.6 ±76.2 sec; Encounter 2: 156.9±41.86 sec) b) 15 minutes of MK-801 (Encounter 1: 574.9±118 sec; Encounter 2: 339.2±122.3 sec). Data is presented as mean±SEM. Asterisks (*) represent significant differences between encounters, with α=0.05.

Similar results were found when the number of bites was evaluated. The second encounter was resolved with lower number of bites than the first one, both when individuals were treated with water (LCL=11.96 bites, UCL=73.4 bites, Z=-2.28, n=10, p-value=0.011, figure 5a) or with MK-801 for 15 minutes (LCL=39.9 bites, UCL=133.9 bites, Z=-3.04, n=10, p-value=0.001, figure 5b).

**Figure 5.**
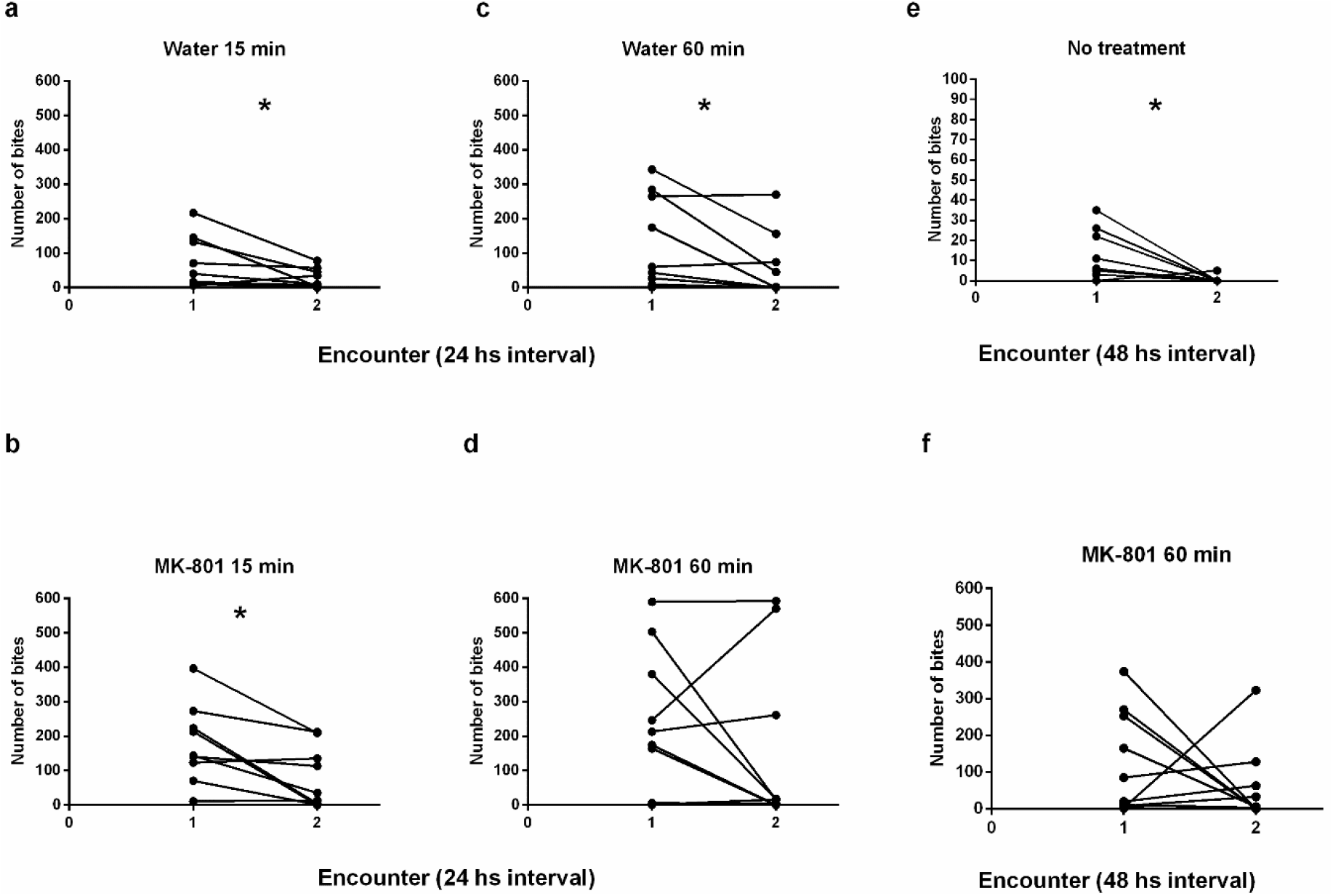
Number of bites performed in the first and second encounter. Each line represents a pair of opponents. Data is presented as mean±SEM. Immediately after the first encounter individuals were exposed to: a) 15 minutes of water and a 24-hour interval between encounters (Encounter 1: 66.6±23.3 bites; Encounter 2: 23.9±8.8 bite). B) 15 minutes of MK-801 and a 24 -hour interval between encounters (Encounter 1: 176.8±38.2 bites; Encounter 2: 80.2±29.6 bites). C) 60 minutes of water and a 24-hour interval between encounters (Encounter 1: 121.2±41.9 bites; Encounter 2: 54.7±28.8 bites). D) 60 minutes of MK-801 and a 24-hour interval between encounters (Encounter 1: 228.1±65.6 bites; Encounter 2: 147.9±76.4 bites). E) No treatment and a 48-hour interval between encounters (Encounter 1: 11.4±3.8 bites; Encounter 2: 0.5±0.5 bites). F) 60 minutes of MK-801 and a 48-hour interval between encounters (Encounter 1: 119.1±43.4 bites; Encounter 2: 55.6±32.4 bites). Asterisks (*) represent significant differences between encounters, with α=0.05.

Given the fact that no effect was found with a 15-minute exposure to MK-801, we designed the second series of experiments using the more prolonged treatment of 60 minutes (Swain et al., 2004). Previously, we performed a pilot experiment in which we evaluate different times (30-, 45- and 60-minutes treatment). Given the fact that we did not find any effect with 30 and 45 min, we chose 60 min for the following experiments.

### 3.4 Effect of 60 minutes of MK-801 exposure in a 24-hour interval

Individuals exposed to 60 minutes of water showed a decrease in the total time of aggressive behaviors in the second encounter, being the difference between days different from zero (LCL=143.21 sec, UCL=294.06 sec, Z=-4.76, n=10, p-value=9.29×10^-7^,Figure 6a) while fish exposed to 60 minutes of MK-801 showed no differences in levels of aggression between encounter 1 and 2 (LCL=-168.46 sec, UCL=522.34 sec, Z=-0.84, n=10, p-value= 0.1997, Figure 6b).

**Figure 6.**
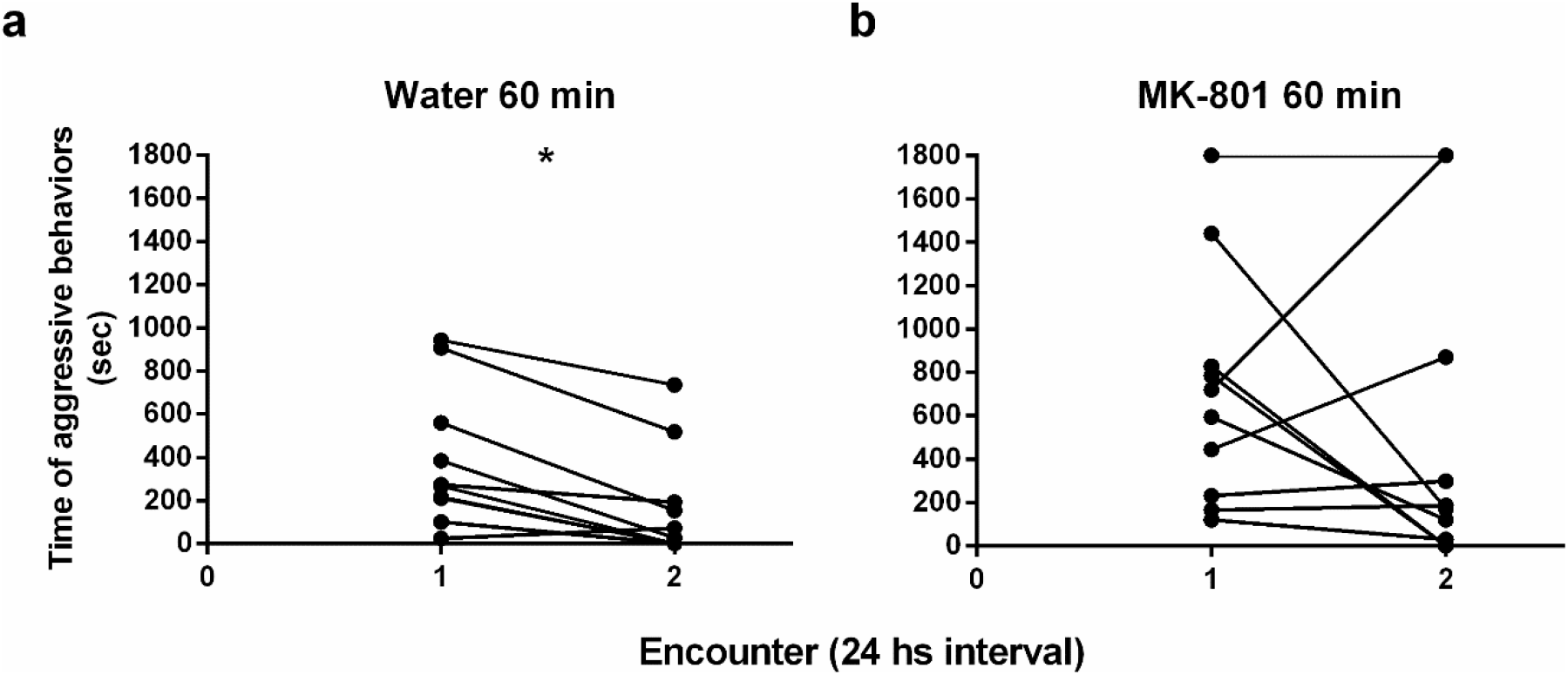
Total time that opponents performed aggressive behaviors (in seconds). Each line represents a pair of opponents with a 24-hour interval between encounters. Immediately after the first encounter individuals were exposed to: a) 60 minutes of water (Encounter 1: 388.9±100.5 sec; Encounter 2: 169.8±80.8 sec.) b) 60 minutes of MK-801 (Encounter 1: 713.1±173.1 sec; Encounter 2: 528±226.5 sec.). Data is presented as mean±SEM. Asterisks (*) represent significant differences between encounters, with α=0.05

When the number of bites was analyzed, the same results of the total time of aggressive behaviors were found for water exposure (LCL=17.09 bites, UCL=115.78 bites, Z=-2.21, n=10, p-value=0.013, figure 5c) and MK-801 exposure (LCL=-38.71 bites, UCL=198.28 bites, Z=-1.10, n=10, p-value=0.134, Figure 5d)

Statistical differences were found in water treatment showing a decrease in aggression in the subsequent encounter, while in the 60 minutes MK-801 treatment no differences were found between days. In all the water experiments the winner of the first encounter won the second one. On the other hand, in MK-801 treatment, in one out of ten final encounters, the winner of the first encounter lost the second one. Furthermore, in this treatment, 7 experiments out of 17 were discarded because on the second day the individuals showed no social interaction possibly due to an effect of the longer treatment with MK-801. To overcome this effect, we used a 48-hours interval between the MK 801 treatment and test. As we shown in the first series of experiments the lower aggression is maintained at least for 72 hours (Figure 2, Figure 3).Therefore, we treated fish with 60 minutes of MK-801, but with a 48-hour interval between fights, in order to avoid any possible unspecific behavioral impact of the drug.

### 3.5 Effect of 60 minutes of MK-801 exposure in a 48-hour interval

When individuals fought and then fought again 48 hours later, the second encounter was resolved with lower levels of aggression, being the difference between days different from zero (LCL=84.33 sec, UCL= 321.73 sec, Z=-2.81, n=10, p-value=0.0024, Figure 7a). Meanwhile, individuals treated with 60 minutes of MK-801 that fought again 48 hours later did not decrease their aggression levels (LCL=-166.99 sec, UCL=412.15 sec, Z=-0.69, n=10, p-value=0.2431, Figure 7b).

**Figure 7.**
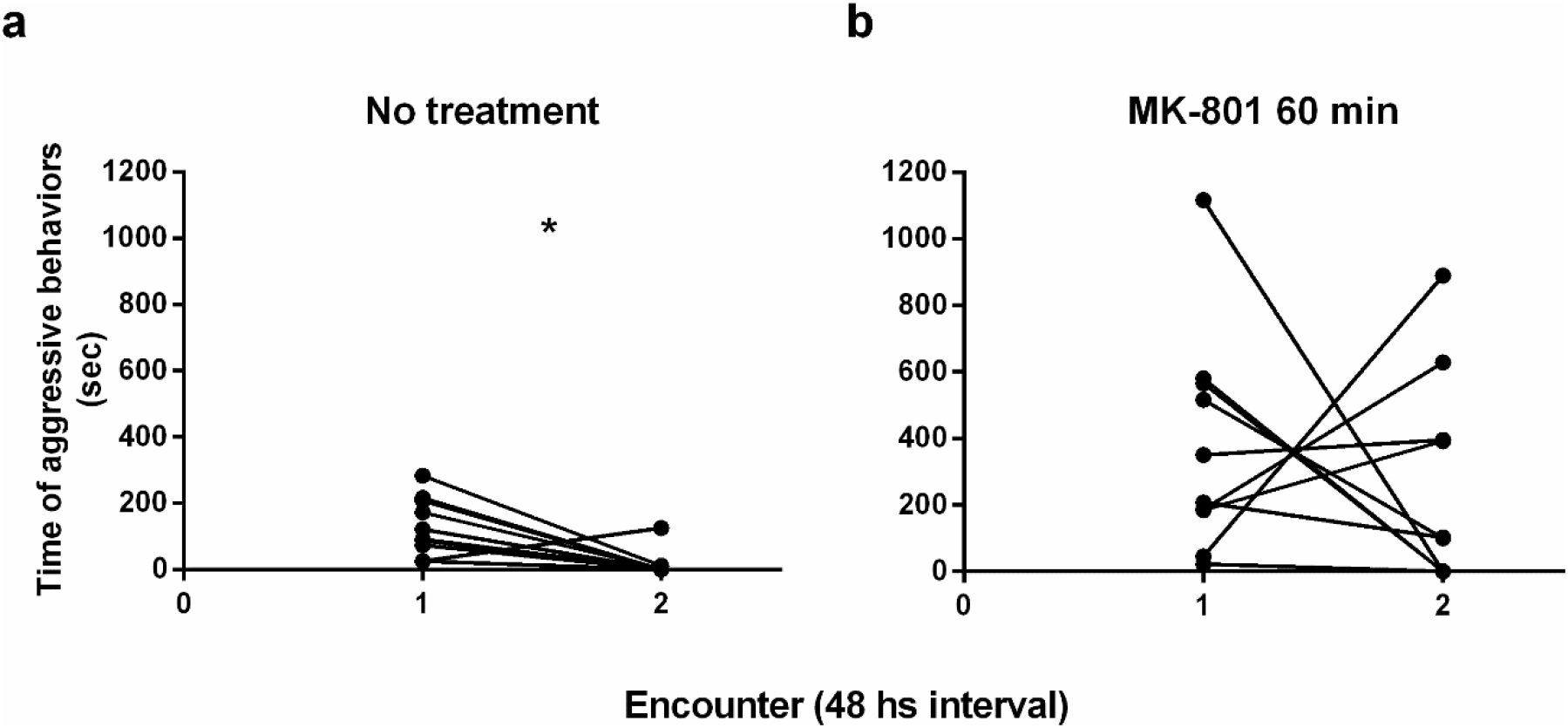
Total time that opponents performed aggressive behaviors (in seconds). Each line represents a pair of opponents with a 48-hour interval between encounters. Immediately after the first encounter individuals were exposed to: a) No treatment (Encounter 1: 179±51.7 sec; Encounter 2: 15.7±12.2 sec.) b) 60 minutes of MK-801 (377.1±104.4 sec; Encounter 2: 250.6±99.2 sec). Data is presented as mean±SEM. Asterisks (*) represent significant differences between encounters, with α=0.05.

Similarly, number of bites decreased in the second encounter (LCL=4.3 bites, UCL= 17.4 bites, Z=-2.71, n=10, p-value=0.003, figure 5e), but this effect was not observed when individuals were treated with MK-801 (LCL=-41.7 bites, UCL=168.2 bites, Z=-0.99, n=10, p-value=0.161) (Figure 5f).

With a 60 minutes exposure of MK-801 and a 48-hour interval, no encounters were discarded and the individuals showed the typical aggressive behaviors previously observed. In all the experiments the winner of the first encounter won the second one.

As a complement to the Bayesian analysis, a frequentist non-parametric analysis (Wilcoxon signed rank test) was performed, showing the same significance of the bayesian analysis (Supplementary material, Table S1, Figure S1, S2).

## 4. Discussion

The present results show that male zebrafish resolve subsequent encounters facing the same opponent with lower levels of aggression, and this effect can be observed within a 72-hour interval between fights. Furthermore, individuals treated with a non-competitive NMDA receptor antagonist (MK-801) immediately after the first encounter do not show this reduction in aggressive behaviors in the second encounter. Therefore, not only the duration of this effect but also its impairment when MK-801 was immediately administered after the first encounter, suggest that the recognition of the opponent, and/or the retrieval of the dynamics and resolution of previous fight may depend on long-term memory.

Memory related to social interaction and individual recognition is very important for individual fitness and a lot of species exhibit this kind of memory. In mice, social memory related to individual recognition is observed after 30 minutes, 24 hours, 3 days, and 7 days between the initial interaction and the test trial (Kogan *et al*., 2000). In laboratory rats, individuals remember and recognize familiar odors after 1- and 48-hour intervals, but not after 96 hours (Burman & Mendl, 2006). Ravens respond differently to familiar and non-familiar playback call after a 3-year separation (Boeckle & Bugnyar, 2012). In other species, memory can last for decades. For example, dolphins can remember and respond to a familiar call after more than a 15-year separation (Bruck *et al*., 2013).

Besides recognizing and remembering each other, individuals can also remember a previous interaction, and change their subsequent behavior. For instance, after hierarchical relationships are established in *Drosophila melanogaster*, the winner retreated less and lunged more, and not only did the opposite happen for the loser, but they also used low-intensity strategies in later fights (Yurkovic *et al*., 2001). In Mangrove rivulus fish, the losing experience increases the probability of immediately retreating from a second encounter and decreases the probability of initiating a new confrontation (Hsu & Wolf, 2001). In these cases, the possibility of remembering an opponent and resolving subsequent encounters with a lower level of aggression can be beneficial for the individuals involved, because they could prevent injuries and wasting energy. Taking this into account, it is expected for this kind of memory to be robust and last for at least 24 hours, although its duration can vary across species.

Paull and coworkers found that male zebrafish establish and maintain hierarchical dominance for at least 5 days in non-interrupted social interactions (Paull *et al*., 2010), which shows that in this species dominance hierarchy lasts several days. In this way, it is expected that if two individuals fight and are then isolated, they would remember their opponent and the previous result and/or the features of the fight for a few days. Our results are in line with this observation.

A previous study conducted in our lab suggests that zebrafish males show a significant decrease in their levels of aggression when faced with the same opponent in a subsequent encounter after a 24-hour interval (Cavallino *et al*., 2020). These results are in concordance with Miklósi and coworkers (Miklósi *et al*., 1992), who have found that paradise fish remember an opponent and change their behavior in a subsequent encounter against the same opponent after a 24-hour interval, but this effect disappears 7 days after the first encounter. In the present work, the behavioral changes in zebrafish males persist for 48 and 72 hours. In both cases, the second encounter is resolved with a lower number of bites and a lower duration of aggression. Further experiments would be necessary to establish the maximum time interval between encounters in which fish remember their opponent and the features of the previous fight.

The recall of a particular opponent and the outcome and features of the first fight allowed zebrafish males to change their subsequent fighting strategies in a 24-, 48- and 72-hour interval between encounters, which suggests the formation of LTM. In this way, the role and mechanism of LTM formation were assessed by the treatment with a non-competitive antagonist of the NMDA receptor.

NMDA receptors are involved in long-term potentiation and long-term memory formation. Consequently, it is well established that inhibiting these receptors has an amnesic effect, impairing LTM (Lutzu & Castillo, 2021). MK-801 is a non-competitive antagonist of this receptor which can be dissolved in water and incorporated by fish. Thus, Blank and coworkers have found that in a single trial inhibitory avoidance, zebrafish show significant retention in the test session, but the exposure of 15 minutes to MK-801 20 μM dissolved in water prevents memory formation (Blank *et al*., 2009). In another associative learning, MK-801 proves to have the same amnesic effect. Zebrafish exposed to a plus maze apparatus can associate a neutral stimulus with the presence of conspecific and remember that learning. Nevertheless, 30 minutes of exposure to MK-801 20 μM impairs memory performance (Sison & Gerlai, 2011). In our work, we have found that 15 minutes of exposure to MK-801 solution is not enough to prevent LTM formation related to an agonistic encounter. This is possibly due to different characteristics between social memory formation and the associative paradigms evaluated in these reports, since the former involves other behavioral and social interactions, and the later are specific associations between other types of stimuli. In sum, only 15 minutes of MK-801 exposure could be not enough to prevent social memory formation in our experiment but prevents memory related to associative learning and inhibitory avoidance tasks. However, when we increase the exposure to 60 minutes, we reveal the same amnesic effect previously reported in other memory paradigms in zebrafish. In a previous work we have demonstrated that after a first dyadic male encounter, when fish are exposed the following day to a second fight against a novel opponent but with the same social status as the original opponent (e.g. always the previous winner versus the previous loser), no reduction in aggressive behaviors is observed in the resolution of the second encounter (Cavallino *et al*., 2020). Similarly, in the present work, when individuals have been treated with an amnesic drug after the first encounter, results show no reduction in aggression during the second encounter.

Exposure to MK-801 has been shown to affect certain behaviors in zebrafish. Swain and coworkers have found that zebrafish treated with 2 and 20 μM MK-801 solutions for 1 hour increase their swimming in circles, but with the 20 μM concentration, the swimming activity does not change (Swain *et al*., 2004). In other reports, this concentration and time exposure has produced motor hyperactivity, increasing the distance traveled and the mean velocity when individuals are tested immediately after the treatment (Seibt *et al*., 2010). Furthermore, treatment with 5 μM solution for 15 minutes induce a decrease in time spent near a mirror and to the segment closest to conspecific, which would be related to a social deficit, and a decrease of aggressive behaviors (Zimmermann *et al*., 2016). A similar social deficit is found in individuals exposed to longer treatments (Perdikaris & Dermon, 2022). In our work, although the second encounter takes place 24 hours after the drug treatment, with a 60-minute exposure some pairs of zebrafish exhibit a deficit in their social interaction during the second encounter. While some individuals show the typical aggressive behaviors, others (possibly more sensitive to the treatment) show no aggressive interactions during the second encounter, exposing a behavioral effect in addition to the effect on memory. However, when the second encounter takes place 48 hours after the drug treatment (and the first fight) all the pairs fight and display typical aggressive behaviors, which shows the effect of impairing memory without the unspecific social deficit observed after 24 hours.

This study suggests first that the lower level of aggression in the second encounter is maintained for at least 72 hours interval between encounters. Secondly, the exposure of zebrafish males to MK-801 for 60 minutes immediately after a fight may prevent the long-term memory formation reflects as behavioral changes observed in the second fight. This behavioral differences may be similar to the effect of changing the first opponent with an unknown one for the second encounter. In both cases the results could be related to remembering an opponent and/or the features and outcome of a previous encounter (Cavallino *et al*., 2020).

## Data availability

Raw video files and data that support these results are available from the corresponding authors upon reasonable request.

## Acknowledgements

To Matias Pandolfi for his contribution to this work, for starting this project and for all these years. To Jimena Santos and Lucia Fernandez Goya for their comments and suggestions to improve the manuscript.

## Supplementary material

**Supplementary Figure 1.**
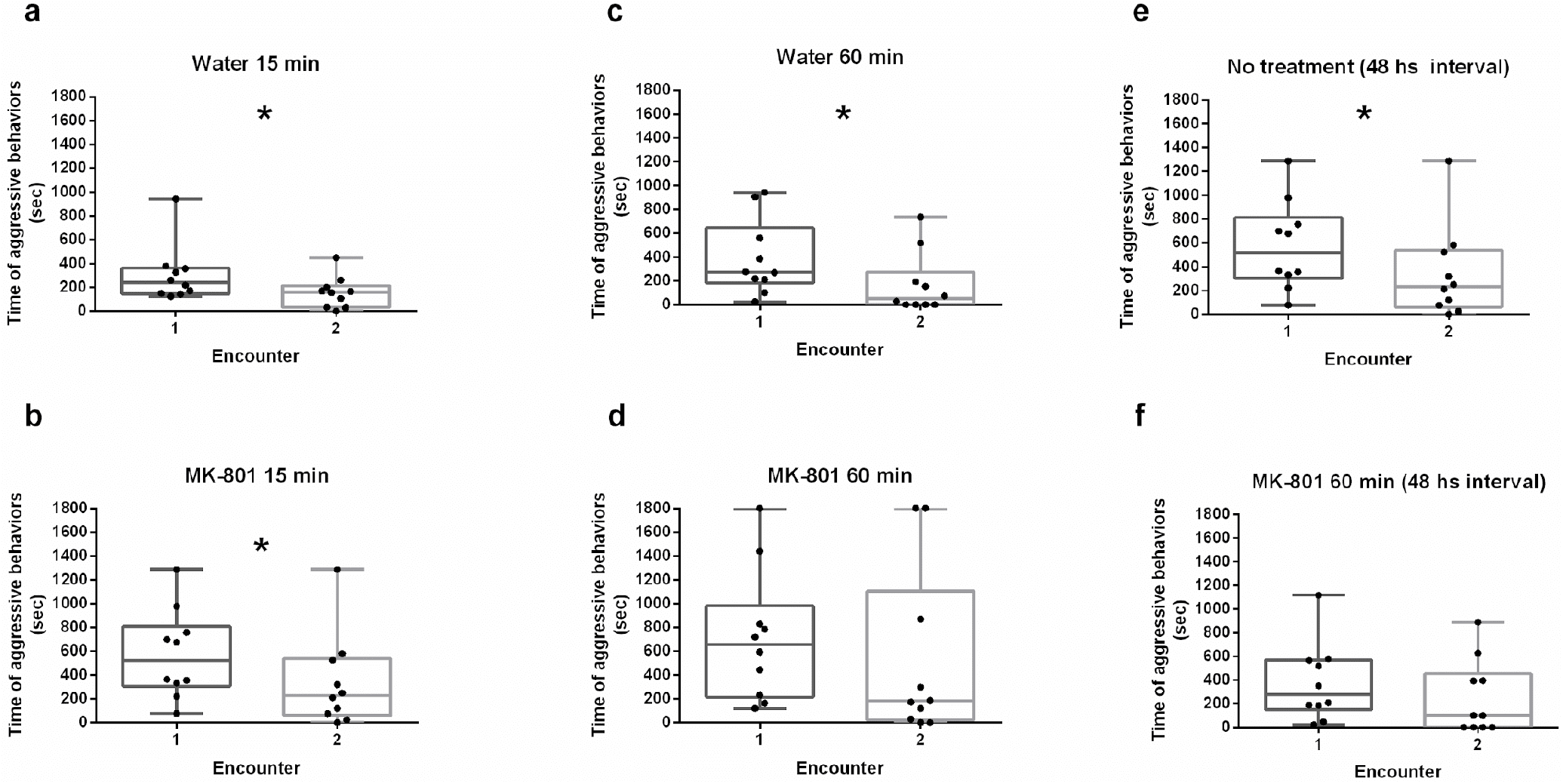
Total time of aggressive behaviors for each treatment. Data is presented as median in seconds and IQR. Band inside the box represents the median and whiskers are the maximum and minimum values. Circles represent the value of each test. Experiments performed with a 24-hour interval where treatment was: a)15 min of water (Encounter 1: 239.5 sec, IQR=195.2; Encounter 2: 161.5 sec, IQR=143.2; N=10; p-value= 0.003906.); b) 15 min of MK-801 (Encounter 1: 520 sec, IQR=404.1; Encounter 2: 230 sec, IQR=386.7; N=10; p-value=0.02439); c) 60 min of water (Encounter 1: 270.5 sec, IQR=303.5; Encounter 2: 49.5 sec, IQR=182.2; N=10; p-value=0.0039); d) 60 min of MK-801 (Encounter 1: 657 sec, IQR=532.7; Encounter 2: 180 sec, IQR=675.5; N=10; p-value=0.2548). Experiments performed with a 48-hour interval where treatment was: e) No treatment (Encounter 1: 106 sec, IQR=121.7; Encounter 2: 0 sec, IQR=2.7; N=10; p-value=0.01953); f) 60 min of MK-801 (Encounter 1: 278.5 sec, IQR=368.2; Encounter 2: 101 sec, IQR=394.2; N=10; p-value=0.5566). Asterisks (*) represent significant differences between encounters, with α=0.05

**Supplementary figure 2.**
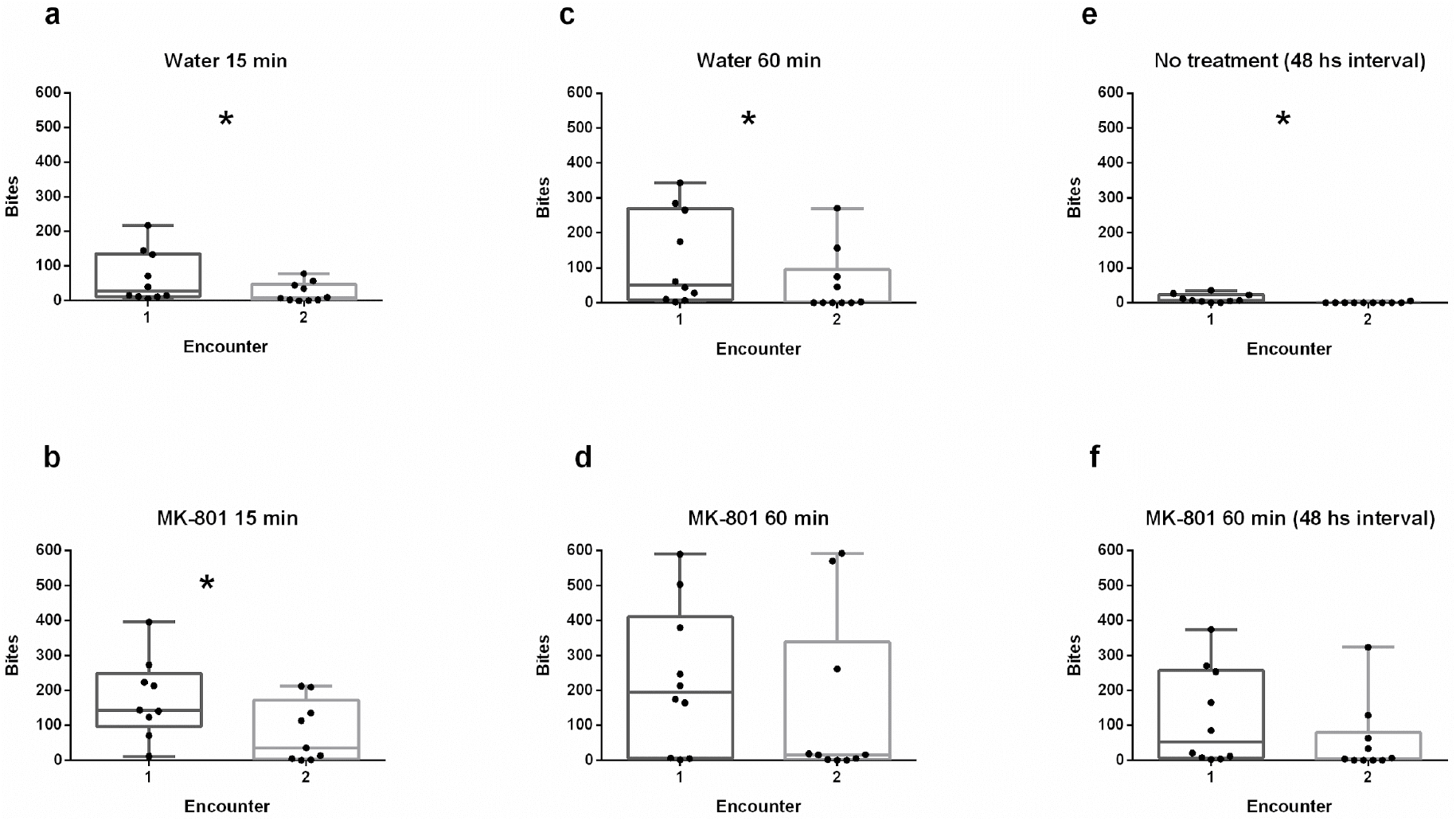
Number of bites for each encounter. Data is presented as median in number of bites and IQR. Band inside the box represents the median and whiskers are the maximum and minimum values. Circles represent the value of each test. Experiments performed with a 24 -hours interval where treatment was: a) 15 min of water (Encounter 1:28 bites, IQR=195.2; Encounter 2: 8.5 bites, IQR=143.2; N=10; p-value=0.039); b) 15 min of MK-801 (Encounter1:143 bites, IQR=100; Encounter 2: 35 bites, IQR=131; N=10; p-value=0.01953). C) 60 minutes of water (Encounter 1: 51.5 bites, IQR=303.5; Encounter 2: 1 bites, IQR=182.2; N=10; p-value=0.048). d) 60 minutes of MK-801 (Encounter 1: 193.5 bites, IQR=532.7; Encounter 2: 15.5 bites, IQR=675.5; N=10; p-value=0.4922). Experiments performed with a 48 - hours interval where treatment was: e) No treatment (Encounter 1: 6 bites, IQR=121.7; Encounter 2: 0 bites, IQR=2.75; N=10; p-value=0.0019); f) 60 min of MK-801 (Encounter 1: 52.5 bites, IQR=368.2; Encounter 2: 4.5 bites, IQR=394.2, N=10; p-value=0.1074). Asterisks (*) represent significant differences between encounters, with α=0.05

**Supplementary Table S1.** P-values of the Wilcoxon signed rank test for all treatments performed.

| Aggression time |  |  | Bites |  |  |
| --- | --- | --- | --- | --- | --- |
| 24 hours interval | 15 min water | 15 min MK | 24 hours interval | 15 min water | 15 min MK |
|  | 0.003906 | 0.02439 |  | 0.03998 | 0.01953 |
| 24 hours interval | 60 min water | 60 min MK | 24 hours interval | 60 min water | 60 min MK |
|  | 0.003906 | 0.2548 |  | 0.04883 | 0.4922 |
| 48 hours interval | No treatment | 60 min MK | 48 hours interval | No treatment | 60 min MK |
|  | 0.01953 | 0.5566 |  | 0.001905 | 0.1074 |

## References

1) Aigner, T. G. Pharmacology of memory: cholinergic—glutamatergic interactions. Curr. Opin. Neurobiol, 5(2), 155–160 (1995).

2) Avdesh, et al. “Regular care and maintenance of a zebrafish (Danio rerio) laboratory: an introduction.” JoVE 69, e4196 (2012).

3) Blank, M., Guerim, L. D., Cordeiro, R. F., & Vianna, M. R. A one-trial inhibitory avoidance task to zebrafish: rapid acquisition of an NMDA-dependent long-term memory. Neurobiol. Learn. Mem, 92(4), 529–534 (2009).

4) Boeckle, M., & Bugnyar, T. Long-term memory for affiliates in ravens. Cur Biol, 22(9), 801–806 (2012).

5) Bruck, J. N. Decades-long social memory in bottlenose dolphins. Proc. Royal Soc. B, 280(1768), 20131726 (2013).

6) Burman, O. H. P., & Mendl, M. Long-term social memory in the laboratory rat (Rattus norvegicus). Anim Welf (2006).

7) Cavallino, L., Dramis, A., Pedreira, M. E., & Pandolfi, M. Effect of previous fighting on the dynamic of agonistic encounters in zebrafish males. Anim. Cogn, 23(5), 999–1006 (2020).

8) Cognato, G. D. P., Bortolotto, J. W., Blazina, A. R., Christoff, R. R., Lara, D. R., Vianna, M. R., & Bonan, C. D. Y-Maze memory task in zebrafish (Danio rerio): the role of glutamatergic and cholinergic systems on the acquisition and consolidation periods. Neurobiol. Learn. Mem, 98(4), 321–328 (2012).

9) Cowan, N. What are the differences between long-term, short-term, and working memory?. Prog brain res, 169, 323–338 (2008).

10) Franscescon, F., Müller, T. E., Bertoncello, K. T., & Rosemberg, D. B. Neuroprotective role of taurine on MK-801-induced memory impairment and hyperlocomotion in zebrafish. Neurochem. Int., 135, 104710 (2020).

11) Gerlai, R. Learning and memory in zebrafish (Danio rerio). Methods Cell Biol. Academic Press 134, 551–586 (2016).

12) Hsu, Y., & Wolf, L. L. The winner and loser effect: what fighting behaviours are influenced?. Anim Behav, 61(4), 777–786 (2001).

13) Huntingford, F. Animal conflict. Springer Science & Business Media (2013).

14) Jarome, T. J., & Helmstetter, F. J. Protein degradation and protein synthesis in long-term memory formation. Front. Mol. Neurosci, 7, 61 (2014).

15) Johnsson, J. I. Individual recognition affects aggression and dominance relations in rainbow trout, Oncorhynchus mykiss. Ethol, 103(4), 267–282 (1997).

16) Kalueff, A. V., Gebhardt, M., Stewart, A. M., Cachat, J. M., Brimmer, M., Chawla, J. et al & Schneider, and the Zebrafish Neuroscience Research Consortium, H. Towards a comprehensive catalog of zebrafish behavior 1.0 and beyond. Zebrafish, 10(1), 70–86 (2013).

17) Kenney, J. W., Scott, I. C., Josselyn, S. A., & Frankland, P. W. Contextual fear conditioning in zebrafish. Learn. Mem, 24(10), 516–523 (2017).

18) Kogan, J. H., Franklandand, P. W., & Silva, A. J. Long-term memory underlying hippocampus-dependent social recognition in mice. Hippocampus, 10(1), 47–56 (2000).

19) Lüscher, C., & Malenka, R. C. (2012). NMDA receptor-dependent long-term potentiation and long-term depression (LTP/LTD). Cold Spring Harb. Perspect. Biol 4(6), a005710.

20) Lutzu, S., & Castillo, P. E. Modulation of NMDA receptors by G-protein-coupled receptors: role in synaptic transmission, plasticity and beyond. Neuroscience, 456, 27–42 (2021).

21) Madeira, N., & Oliveira, R. F. Long-term social recognition memory in zebrafish. Zebrafish, 14(4), 305–310 (2017).

22) McGaugh, J. L., & Izquierdo, I. The contribution of pharmacology to research on the mechanisms of memory formation. Trends Pharmacol. Sci 21(6), 208–210 (2000

23) McGaugh, J. L. Memory--a century of consolidation. Science, 287(5451), 248–251 (2000).

24) Miklósi, Á., Haller, J., & Csányi, V. Different duration of memory for conspecific and heterospecific fish in the paradise fish (Macropodus opercularis L.). Ethol, 90(1), 29–36 (1992).

25) Mourier, J., Brown, C., & Planes, S. Learning and robustness to catch-and-release fishing in a shark social network. Biol. Lett., 13(3), 20160824 (2017).

26) Nicoll, R. A., & Malenka, R. C. Expression mechanisms underlying NMDA receptor-dependent long-term potentiation. Ann. N. Y. Acad. Sci., 868(1), 515–525 (1999).

27) Oliveira, R. F., Silva, J. F., & Simoes, J. M. Fighting zebrafish: characterization of aggressive behavior and winner–loser effects. Zebrafish, 8(2), 73–81 (2011).

28) Paull, G. C., Filby, A. L., Giddins, H. G., Coe, T. S., Hamilton, P. B., & Tyler, C. R. Dominance hierarchies in zebrafish (Danio rerio) and their relationship with reproductive success. Zebrafish, 7(1), 109–117 (2010).

29) Perdikaris, P., & Dermon, C. R. (2022). Behavioral and neurochemical profile of MK-801 adult zebrafish model: Forebrain β2-adrenoceptors contribute to social withdrawal and anxiety-like behavior. Progress in Neuro-Psychopharmacology and Biological Psychiatry, 115, 110494

30) Scaia, M. F., Akinrinade, I., Petri, G., & Oliveira, R. F. Sex differences in aggression are paralleled by differential activation of the brain social decision-making network in zebrafish. Front behav neuro, 16 (2022).

31) Seibt, K. J., da Luz Oliveira, R., Zimmermann, F. F., Capiotti, K. M., Bogo, M. R., Ghisleni, G., & Bonan, C. D. Antipsychotic drugs prevent the motor hyperactivity induced by psychotomimetic MK-801 in zebrafish (Danio rerio). Behav brain res, 214(2), 417–422 (2010).

32) Sison, M., & Gerlai, R. Associative learning performance is impaired in zebrafish (Danio rerio) by the NMDA-R antagonist MK-801. Neurobiol. Learn. Mem, 96(2), 230–237 (2011).

33) Swain, H. A., Sigstad, C., & Scalzo, F. M. Effects of dizocilpine (MK-801) on circling behavior, swimming activity, and place preference in zebrafish (Danio rerio). Neurotoxicol Teratol, 26(6), 725–729 (2004).

34) Triki, Z., & Bshary, R. Long-term memory retention in a wild fish species Labroides dimidiatus eleven months after an aversive event. Ethol, 126(3), 372–376 (2020).

35) Yurkovic, A., Wang, O., Basu, A. C., & Kravitz, E. A. Learning and memory associated with aggression in Drosophila melanogaster. Proc. Natl. Acad. Sci, 103(46), 17519–17524 (2006).

36) Zimmermann, F. F., Gaspary, K. V., Siebel, A. M., & Bonan, C. D. Oxytocin reversed MK-801-induced social interaction and aggression deficits in zebrafish. Behav. Brain Res, 311, 368–374 (2016).

